# A large-scale comparative metagenomic analysis of short-read sequencing platforms indicates high taxonomic concordance and functional analysis challenges

**DOI:** 10.1101/2025.07.06.662369

**Authors:** Kinga Zielińska, Kateryna Pantiukh, Paweł P Łabaj, Tomasz Kosciolek, Elin Org

## Abstract

Driven by the increasing scale of microbiome studies and the rise of large, continuously expanding population cohorts, the volume of sequencing data is growing rapidly. As such, ensuring the comparability of data generated across different sequencing platforms has become a pressing concern in efforts to uncover robust links between the microbiome and human health. In this study, we conducted a comprehensive comparison of taxonomic and functional profiles from 1,351 matched human gut microbiome sample pairs, sequenced using both the MGISEQ-2000 (MGI) and NovaSeq 6000 (Illumina NovaSeq) platforms. Taxonomic profiles showed high concordance within and between platforms: 96.44 ± 5.96% of species were shared between MGI–MGI pairs, and 92.07 ± 5.20% were shared between MGI and NovaSeq pairs. The proportion of platform-specific species was low, at 3.42% for MGI–MGI comparisons and 5.89% for MGI–NovaSeq comparisons. No significant differences in Shannon diversity were observed for either within-platform or between-platform comparisons. However, functional profiles revealed notable discrepancies between platforms, which were attributed to differences in pre-sequencing protocols.

**Importance:** Our findings demonstrate robust taxonomic comparability between MGI and NovaSeq platforms, while revealing systematic functional differences that should be carefully considered in cross-platform and even cross-cohort metagenomic studies.

## Introduction

Metagenomics has emerged as a powerful tool for uncovering key roles that microorganisms play in human health and environmental ecosystems (1),(2). Recent advancements in Next Generation Sequencing (NGS) technologies have enabled increasingly fine-scale taxonomic resolution, while also significantly expanding our understanding of microbial functions and their dynamic interactions with hosts and other microbes. However, the expansion of NGS has been followed by new protocols and sequencing platforms. This introduced biases, complicating interpretation and reducing the reproducibility of signals identified through metagenomic analyses.

A recent study by Forry et al. compared the Firmicutes:Bacteroides ratios defined from the same biological samples independently by 44 participating labs (3). The results revealed a substantial impact of a protocol choice, introducing both biases in measurements tied to specific methodological decisions, as well as influencing measurement robustness, as evidenced by the variability in results across labs employing similar methods. Metagenomic results - specifically microbial load and composition - are highly susceptible to biases stemming from sample storage. This is particularly true for low-biomass samples, underscoring the necessity of an efficient, universal storage method to ensure robust cross-study comparisons (4).

The major providers of massive parallel sequencing technologies in metagenomics are Illumina (sequencing by synthesis) and MGI/BGI (DNBSEQ™) technology, both claiming accuracy but differing substantially in underlying chemistry and amplification design. These methodological distinctions can introduce systematic variation in metagenomic data. A key difference lies in the DNA amplification strategy. Illumina’s sequencing-by-synthesis (SBS) technology employs bridge amplification, a solid-phase exponential PCR on a glass flow cell that is known to be susceptible to GC-content bias and the propagation of early-cycle PCR errors (5). In contrast, MGI’s DNBSEQ™ technology utilizes rolling-circle amplification (RCA) to generate DNA Nanoballs (DNBs) from single DNA molecules in solution (6). This linear amplification process minimizes amplification bias and reduces PCR duplication compared with exponential bridge amplification. Additional differences exist in the sequencing chip architecture, where MGI loads DNBs onto high-density patterned arrays, which may reduce optical noise compared to the random cluster distribution on Illumina flow cells (7). Beyond sequencing chemistry, upstream steps such as DNA extraction and library preparation represent further a well-documented source of experimental bias that can influence metagenomic profiles (8). Moreover, differences in multiplexing levels between sequencing runs can influence the relative representation of libraries, leading to systematic variation in sequencing depth and read distribution across samples. Collectively, these technical distinctions are potential sources of systematic variation that can influence the results of metagenomic studies.

Comparative analyses of sequencing by synthesis and DNBSEQ technologies conclude that the two could be used interchangeably. They cover a range of fields, starting with whole-genome DNA sequencing (9), through paleogenomics (10) and bacterial genome assembly. Multiple sequencing-specific biases has been discovered, however, including a higher number of insertions detected in HiSeq 2000 and more deletions in BGISEQ-500 genome assemblies, as well as a bias toward genes with higher GC content on the HiSeq platforms (11). These could have profound impacts on the downstream analyses, but they have not been further explored.

Here, we take advantage of a large population-based metagenomic dataset (1,351 individuals) simultaneously sequenced using the Illumina NovaSeq 6000 (NovaSeq) and MGISEQ-2000 (MGI) platforms, allowing us to directly compare the sequencing technologies for metagenomic applications (12). We begin by evaluating the robustness of MGI sequencing through an analysis of samples sequenced twice (N=53), assessing the degree of intra-sequencing variability. We then conduct a comprehensive comparative analysis of the full dataset, examining matched samples from MGI and NovaSeq platforms to identify sequencing-specific biases (N=1,351).

## Results

### Study design and cohort overview

Our aim was to evaluate the consistency and comparability of microbiome sequencing outputs across the platforms, with a specific focus on the taxonomic and functional profiles. We therefore utilized a subset of the Estonian microbiome cohort (EstMB), sequenced on NovaSeq and MGI platforms.

The original EstMB cohort is a population-based cohort consisting of 2,509 gut metagenomic samples, sequenced on the NovaSeq 6000 platform (hereafter NovaSeq) to 30.63 ± 3.12 million paired reads per sample (**Figure 1a)** (12). A subset of the EstMB cohort was sequenced on the MGISEQ-2000 platform by MGI Tech Co, Ltd (hereafter MGI) with deeper coverage (108.71 ± 42.08 million paired reads per sample). This subset, referred to as the Estonian Microbiome deep cohort (EstMB-deep), includes 1,729 gut metagenomic samples (53 samples sequenced twice), all of which are also represented in the EstMB cohort **(Figure 1a, 1b).**

**Figure 1.**
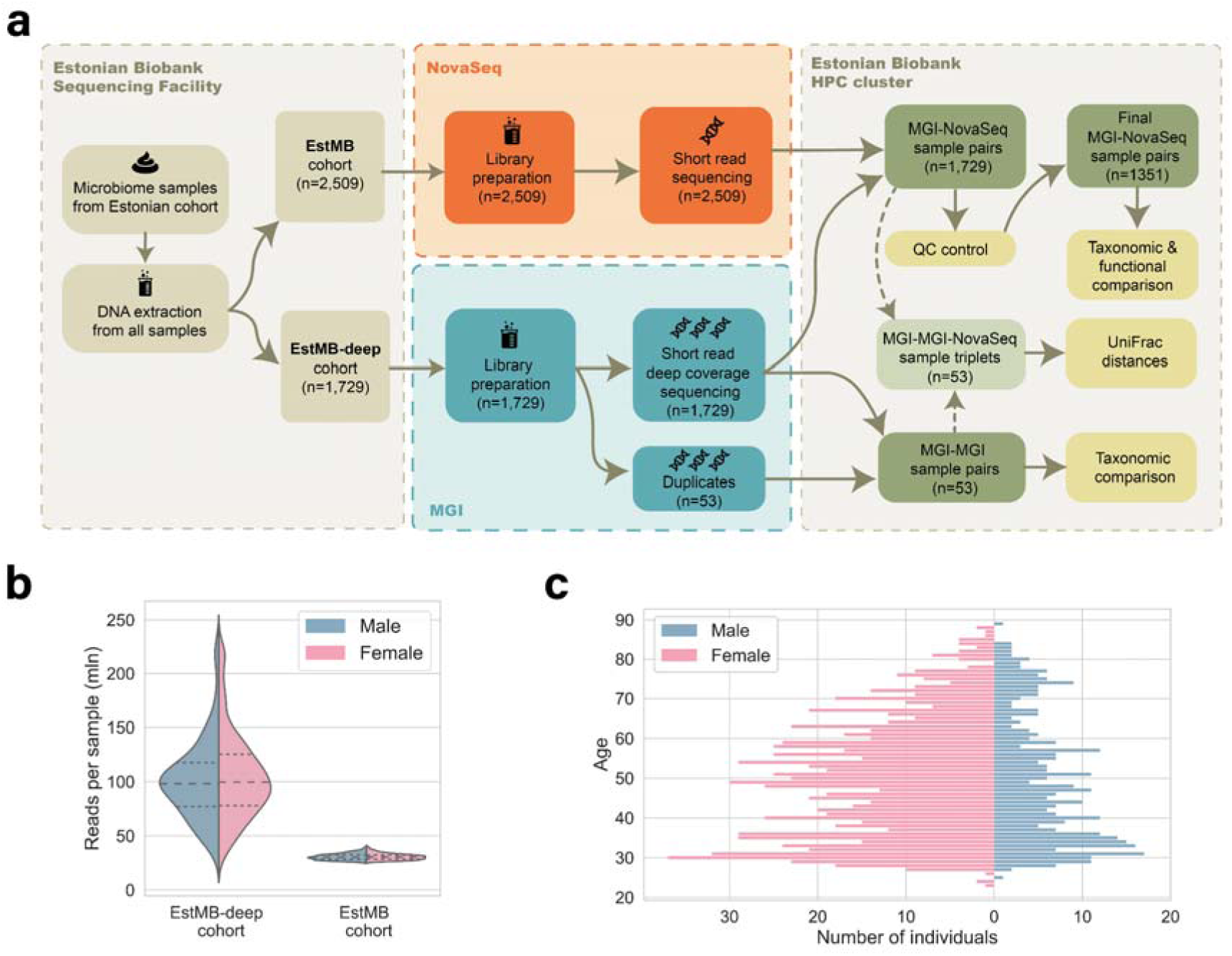
Study Overview. **a.** Study design and cohort characteristics **b.** Distribution of the number of reads per sample for MGI (EstMB-deep cohort) and NovaSeq (EstMB cohort) sequenced samples from the final sample set (N=1,351). **c.** Gender and age distributions of individuals in the final sample set (N=1,351).

To evaluate within-platform and cross-platform variability, we first analysed the MGI–MGI pairs (53 samples randomly selected and sequenced on the MGI platform twice). We then expanded the analysis to include 1,729 samples sequenced on both platforms: MGI–NovaSeq pairs. To ensure a fair comparison between platforms, MGI reads were subsampled first to match the read depth of their NovaSeq counterparts. As a result of the initial quality assessment, the high-quality dataset comprised 1,351 MGI–NovaSeq pairs and consisted of 71.87% females, ranging in age from 23 to 89 years with a mean age of 49.7 years **(Figure 1c).** The samples then underwent downstream profiling and subsequent cross-platform comparisons. (**Figure 1a**).

### Technical reproducibility: intra-platform comparison of MGI replicates

To establish a baseline for platform consistency, we first evaluated intra-platform variability by examining whether replicate sequencing runs introduced systematic variation. Specifically, we analysed 53 biological samples sequenced in duplicate on the MGISEQ-2000 platform within the same library preparation batch. This approach ensured the observed reproducibility reflectd technical consistency of sequencing within a single batch, isolated from library preparation variation. To assess intra-platform taxonomic consistency, we analysed MGI–MGI sample pairs (replicates from MGI-1 and MGI-2) which were rarefied to equal read counts (**Figure 1a**). Consistency was evaluated using metrics comparing the two sets, including the percentage of shared species, the difference in the number of detected species, and the differences in species prevalence, relative abundance, and Shannon diversity index.

Across the two MGI sequencing runs, a total of 1,083 species (380 genera) were identified. Only 3.42% (37 species) were detected exclusively in one run: 17 species unique to MGI-1 and 20 to MGI-2 (**Figure 2a, Supplementary Table S1**). These set-specific species exhibited low prevalence and low relative abundance. Specifically, 36 of the 37 were found in only one sample (one species, *Lactococcus cremoris*, was found in two). The mean relative abundance of set-specific species was 2.25 × 10⁻LJ %, which contrasts with the 7.67 × 10⁻LJ % mean relative abundance for the shared species (**Figure 2b, Supplementary Figure 1**).

**Figure 2.**
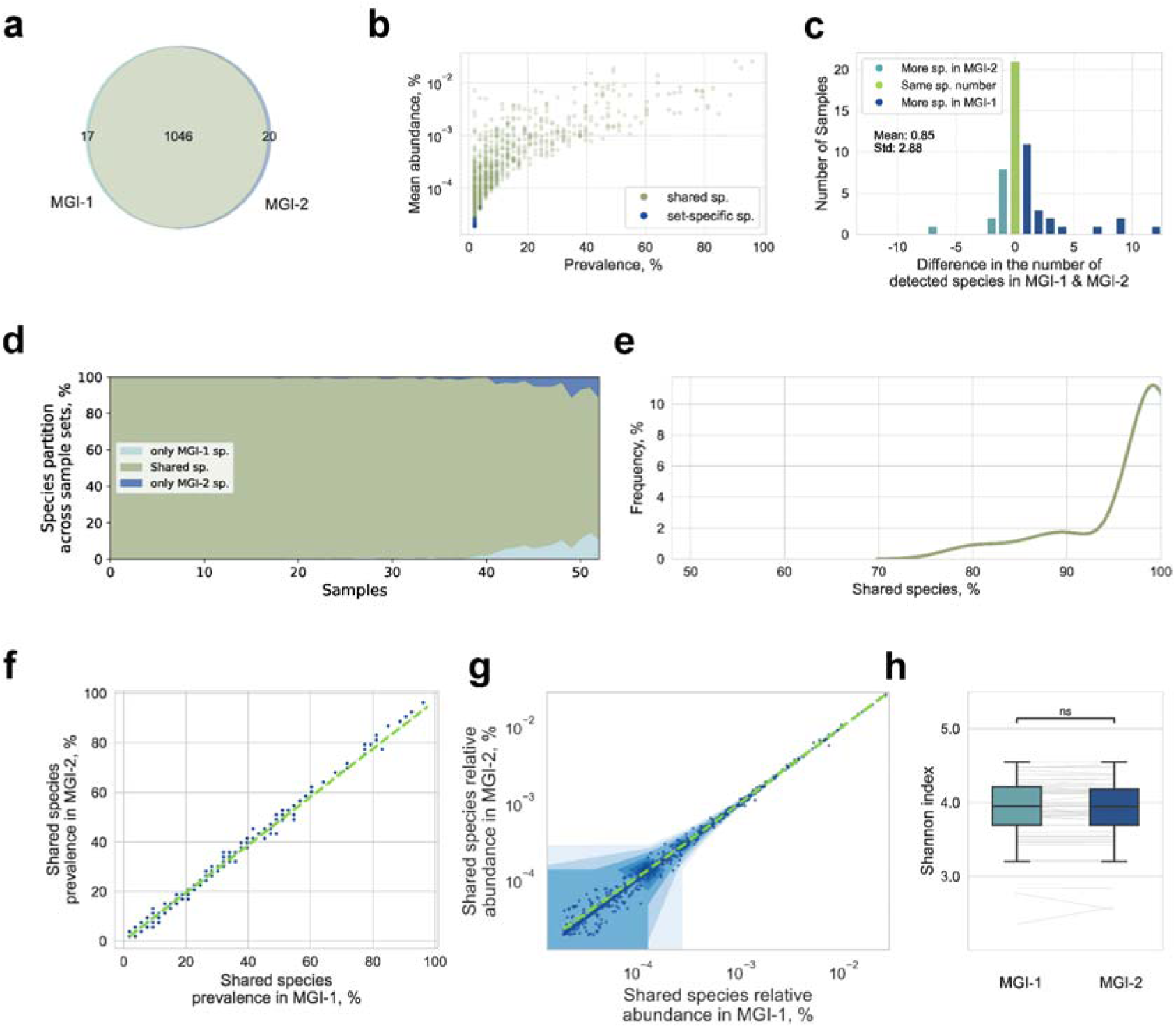
Intra-platform baseline comparison. (N=53). **a.** Venn diagram of the number of species detected in MGI-1 and MGI-2. **b.** Prevalence versus mean relative abundance of detected species, with species found only in one run highlighted in blue. **c.** Distribution of differences in the number of detected species within sample pairs. Pairs with identical species counts are represented by green bars, while sample pairs with more species in MGI-1 or MGI-2 are represented by light or dark blue bars, respectively. **d.** The percentage of shared and unique run-specific species within each sample pair. **e.** Distribution of the percentage of shared species across sample pairs. **f.** Comparison of the prevalence of shared species between matched samples. **g.** Comparison of the relative abundance of shared species in matched samples. **h.** Comparison of the Shannon diversity Index within sample pairs.

To assess the consistency of species detection, we calculated the difference in species counts for each MGI–MGI sample pair and visualized the distribution (**Figure 2c**). The distribution was centered around a mean difference of 0.85 species (SD=2.88). The Wilcoxon signed-rank test showed a significant difference between runs (p=0.032), but with a small effect size (Cohen’s d=0.29). This suggests the observed difference was minor and likely due to random variation (e.g., stochasticity in library preparation, lane/batch effects, or minor read assignment fluctuations) rather than true biological differences.

The overlap of species detected within MGI–MGI sample pairs demonstrated a high level of concordance. Most pairs shared more than 90% of species (mean = 96.44% ± 5.96%, **Figure 2d and 2e**). Furthermore, species prevalence and relative abundance showed strong agreement across sequencing runs, with high correlation indicated by R^2^ values approaching 1.0 (**Figure 2f and 2g**). Finally, Shannon diversity index values were consistent across replicates, with a paired t-test revealing no significant differences (T-statistic = 0.086, p > 0.05, Cohen’s d: 0.017, **Figure 2h**). This indicated consistent diversity estimates across the technical replicates.

### Cross-platform reproducibility of taxonomic composition

Following the assessment of intra-platform variability, we addressed the central question: whether MGI and NovaSeq sequencing platforms produce biologically comparable results. We conducted a comprehensive analysis of paired samples generated on both platforms. This evaluation included a comparison of taxonomic composition, raw sequencing read quality, and functional profiling.

#### Assessment of intra-and inter-platform variability using MGI-MGI-NovaSeq triplets

Prior to the large-scale analysis, we assessed intra- and inter-platform variability using Shannon entropy and UniFrac distances on a subset of 53 samples sequenced three times: twice on MGI and once on NovaSeq (MGI–MGI–NovaSeq triads). We used the UniFrac metric to quantify differences based on phylogenetic relationships. We found no statistically significant difference between intra-platform (MGI vs. MGI) and inter-platform (MGI vs. NovaSeq) UniFrac distances (PERMANOVA p-value=1; **Supplementary Figure 2a**), and matched samples showed substantial overlap in the three-dimensional PCoA space (**Supplementary Figure 2b**). However, in 79% of the triads, the two MGI replicates were more similar to each other than to the corresponding NovaSeq replicate. Furthermore, the mean distance between MGI and NovaSeq samples was approximately 2.5 times greater than the distance between the MGI replicates, demonstrating that intra-platform variability was consistently lower than cross-platform variability.

### Raw reads sequencing quality assessment

To assess raw sequencing quality, we ran FastQC on all samples from the MGI–NovaSeq sample pairs (1,729 samples per platform). All samples passed key quality metrics, including per-base sequence quality and adapter content. One sample from each platform was flagged for “Overrepresented sequences” (those samples were from different pairs). GC content distributions were overall similar between the platforms, with a mean of 45.27D±D1.82% for MGI and 45.60D±D1.61% for NovaSeq. Despite the similarity, a paired Wilcoxon test indicated a statistically significant difference in the GC content (p = 4.25 × 10⁻²⁵), although the effect size was small (Cohen’s d ≈ 0.24) (**Figure 3a**). The slightly higher GC content observed in NovaSeq samples aligns with the platform’s known bias against low-GC regions, likely attributable to PCR amplification and cluster generation artifacts inherent to Illumina’s bridge amplification (13). In contrast, DNBSEQ technology, utilizing rolling circle amplification to generate DNA nanoballs before loading onto a patterned array, aims to minimize such amplification biases.

**Figure 3.**
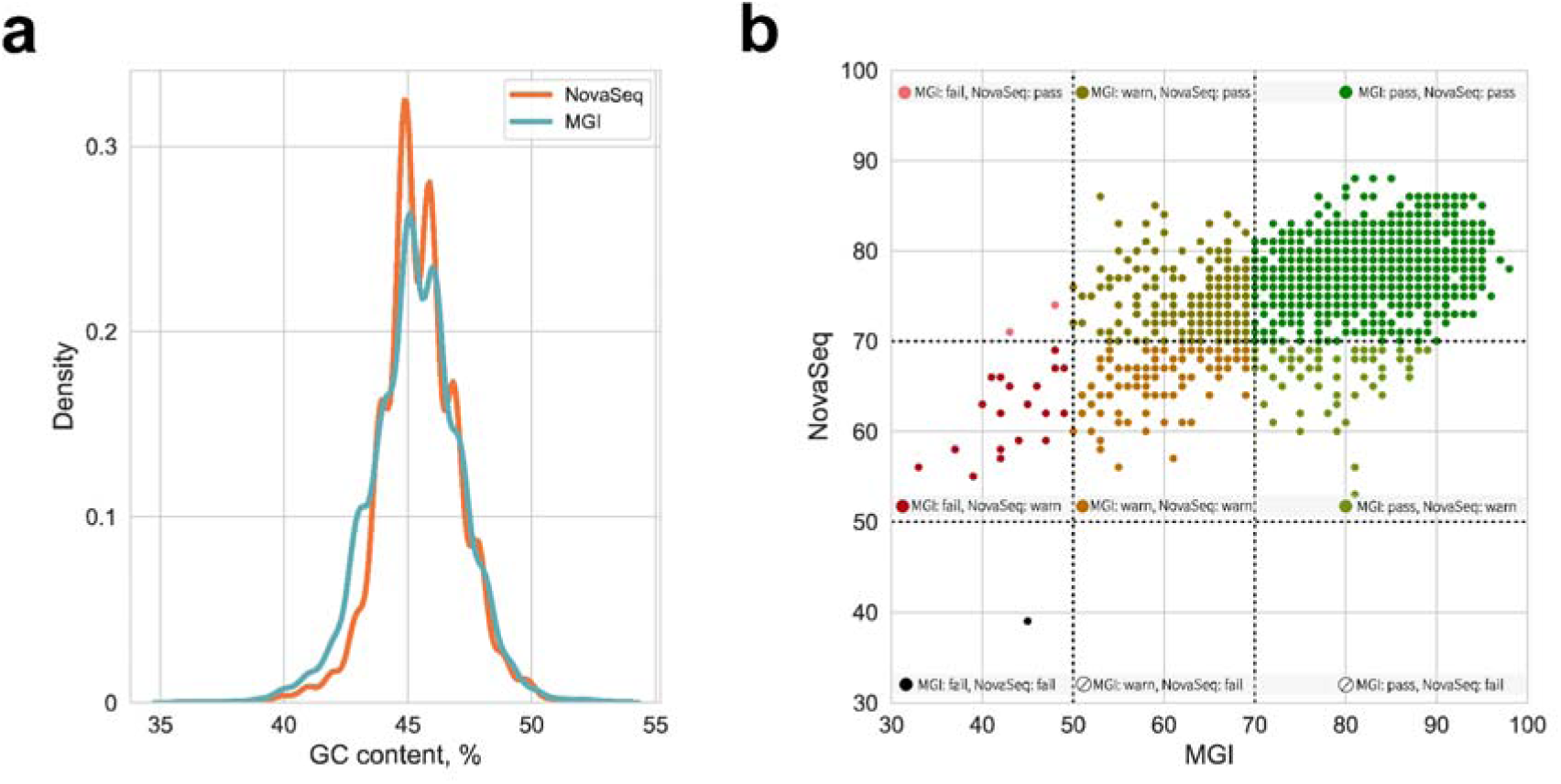
Inter-platform raw sequencing quality comparison (n=1,729) **a.** Distribution of the GC content across MGI and NovaSeq samples **b.** Sample pairs with their corresponding quality tags, assigned based on sequence duplication levels.

Based on sequence duplication levels, 12.2% of all samples did not meet quality thresholds and were flagged as “warn” or “fail”, indicating potential issues such as PCR amplification bias, low library complexity, or contamination. The failure rate was substantially higher for MGI (15.9%) compared to NovaSeq (8.4%) (**Figure 3b**). Notably, the failed samples did not consistently overlap across platforms, suggesting that the observed quality issues likely reflected platform-specific biases, differences in library preparation, or other technical artefacts. If this had been due to early-stage sample processing errors, both platforms would have been affected in a similar manner.

To ensure the robustness of the comparison, sample pairs containing at least one quality warning (“warn” or “fail”, N=378) were excluded from downstream analysis. This yielded a final dataset of 1,351 high-quality sample pairs, which were subsequently used for taxonomic and functional assessments.

### Cross-platform concordance in taxonomic profiles

We performed taxonomic comparisons for all sample pairs from the final dataset (N=1,351). Across both sequencing platforms, we identified a total of 2,953 species spanning 808 genera. Among these, 5.89% (174 species) were detected exclusively by one platform - 88 species were unique to MGI and 86 to NovaSeq (**Figure 4a, Supplementary Table S2**). Most of the platform-specific taxa were rare: 154 out of 174 appeared in only one sample, and their mean relative abundance (1.30 × 10⁻LJ %) was markedly lower than that of the shared species (2.85 × 10⁻LJ %) (**Figure 4b**).

**Figure 4.**
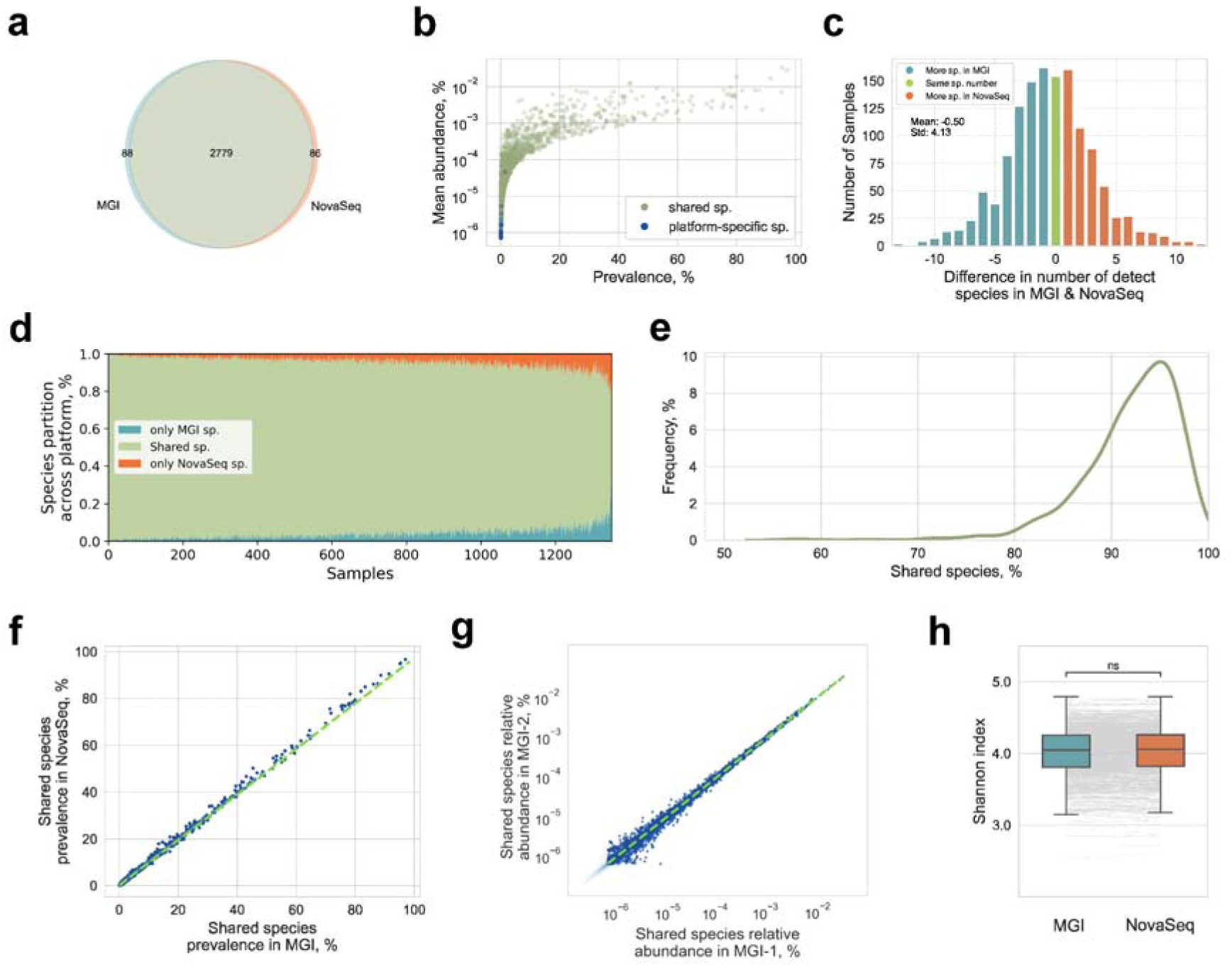
Taxonomic composition comparison between sequencing platforms (n=1,351) a. Proportion of species shared between platforms and those detected exclusively in MGI or NovaSeq. b. Prevalence and relative abundance of all detected species, with platform-specific species (found only in one platform) highlighted in blue. c. Distribution of differences in the number of species detected per sample pairs. Pairs with identical species counts are shown in green, those with more species in MGI are shown in light blue and with more species in NovaSeq are shown in orange. d. Percentage of shared and platform-specific species across sample pairs e. Distribution of the percentage of shared species within each sample pair f. Species prevalence for taxa shared between platforms within sample pairs. g. Relative abundance of shared species within sample pairs h. Comparison of Shannon diversity index within each sample pair.

To assess the consistency in species detection, we compared the difference in species counts for each paired sample and visualized the distribution of these differences (**Figure 4c**). The distribution was centered around a mean difference of -0.50, with a standard deviation of 4.13. The distribution failed the Shapiro–Wilk test (p ≪ 0.05), prompting rejection of its normality and the use of the Wilcoxon signed-rank test, which showed a significant difference in species counts between platforms (p = 1.03 × 10⁻D). However, the effect size was small (Cohen’s d = -0.12), and visual inspection suggested the difference was minimal and likely due to random variation.

To extend our comparison, we assessed the overlap of species detected within the matched samples. As shown in **Figures 4d** and **4e**, most pairs exhibited a high level of concordance, with more than 90% of species shared between the replicates (mean = 92.07 ± 5.20%). In addition, species prevalence and relative abundance exhibited strong agreement across sequence runs, demonstrating high correlation with R^2^ values approaching 1.0 (**Figures 4f** and **4g, Supplementary Figure 3**).

No significant difference in Shannon diversity was observed between the platforms (paired t-test, p > 0.05) (**Figure 4h**). Additionally, Procrustes analysis demonstrated excellent concordance between sample pairs across the two platforms (M² = 0.02, p = 0.0001; **Supplementary Figure 4**), we also identified no statistically significant beta diversity differences (Bray-Curtis and Euclidean distances, PERMANOVA p-values > 0.05). Differential abundance analysis with MaAsLin2 initially revealed three species enriched in either platform (two from the *ER4* genus family *Oscillospiraceae* and one *Paraprevotella*), but they all failed to pass the correction for multiple testing (**Supplementary Table S3**).

To test the biological interpretability of the results, we examined associations between microbial taxa and health-related variables using linear regression, separately for each sequencing platform. Taxonomic associations were overall consistent between the MGI and NovaSeq datasets, with only two taxa showing platform-specific significance but with negligible effect sizes (**Supplementary Table S4**).

The results of the intra- and inter-platform taxonomic comparisons are summarised in **Table 1**.

**Table 1.** Comparison summary: intra-platform (MGI-MGI) and inter-platform (NovaSeq-MGI) taxonomy metrics.

| Metric | Intra-platform comparison (MGI-MGI sample pairs) | Inter-platform comparison (NovaSeq-MGI sample pairs) |
| --- | --- | --- |
| Number of sample pairs | 53 | 1,351 |
| Percentage of set-specific species | 3.42% (37 out of 1,083 total species) | 5.89% (174 out of 2953 total species) |
| Difference in the number of | significant difference | significant difference |
| species per sample pair<br>(Wilcoxon signed-rank test) | (p = 0.038)<br>Effect size: small<br>(Cohen's $d = 0.29$) | (p = $1.03 \times 10^{-7}$)<br>Effect size: very small<br>(Cohen's $d = -0.12$) |
| Mean percentage of shared species within sample pairs | 96.44% ± 5.96% | 92.07% ± 5.20% |
| Correlation of species' prevalence and relative abundance between platforms within sample pairs | $R^2=0.999$ | $R^2=0.999$ |
| Difference in Shannon diversity index between platforms within sample pairs (paired t-test) | p > 0.05 | p > 0.05 |

### Platform-specific functional differences reflect sequencing depth and complexity

To systematically assess differences in functional profiling between the sequencing platforms, we ‘randomly selected 700 matched sample pairs from our final dataset of 1,351 metagenomic samples. Functional annotation was performed with *mi*-faser, which resulted in the identification of 1,170 unique enzymes (EC numbers) across all samples. Of these, 2.65% (31 enzymes) were detected exclusively by one platform - 17 enzymes uniquely to MGI and 14 uniquely to NovaSeq (**Figure 5a** and **b**).

**Figure 5.**
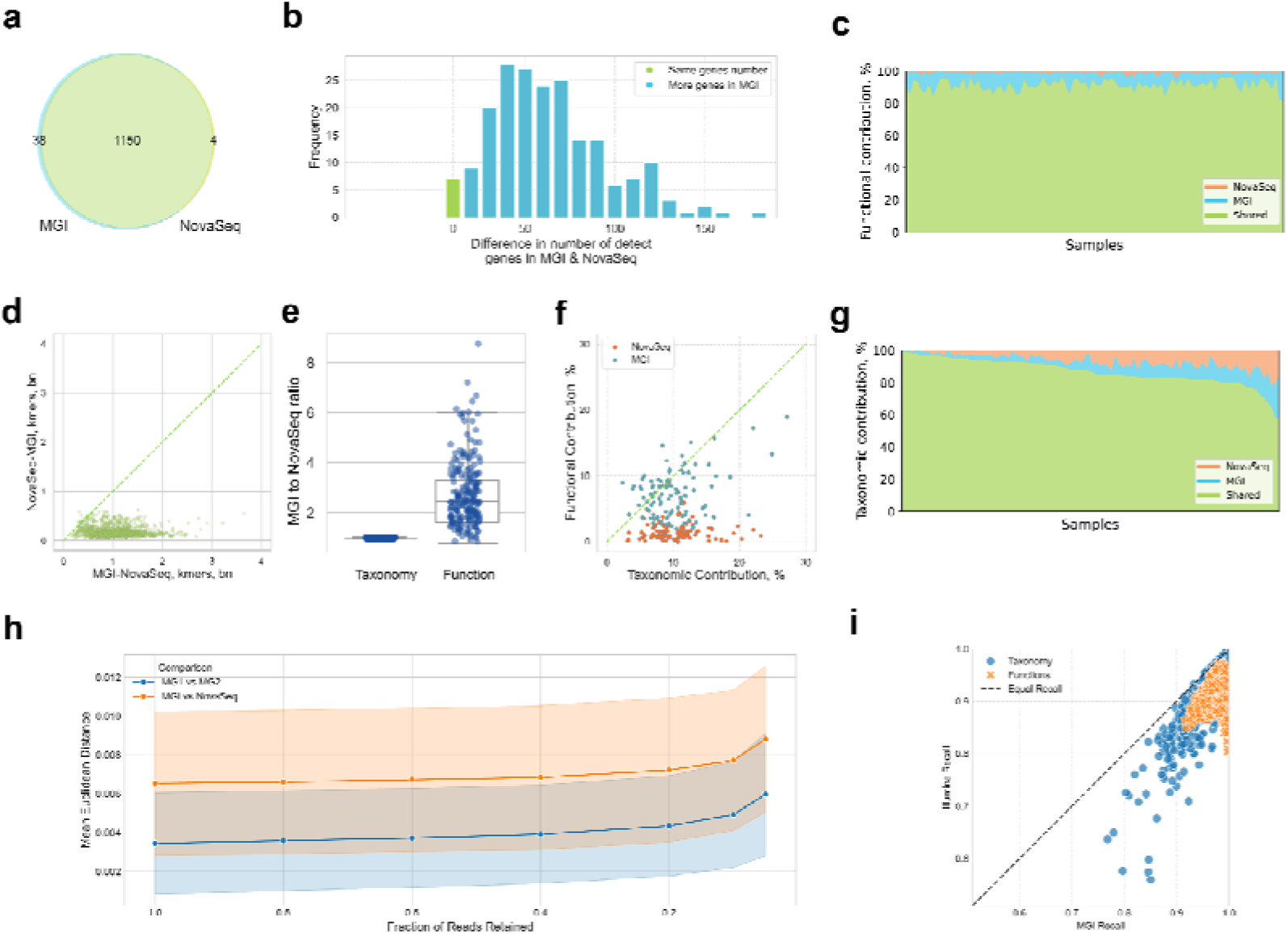
Function-based comparison of the MGI and NovaSeq batches (n=700). (a) Proportion of genes shared between platforms and platform-specific genes, detected exclusively in MGI or NovaSeq platforms. (b) Overlap of functions in matched MGI and NovaSeq samples. c) A functional overlap in 200 exemplary samples with different levels of taxonomic overlap, sorted by descending functional overlap. (d) Unique k-mers per sample: in NovaSeq versus MGI and vice versa. (e) Contributions of taxonomic and functional microbiome components found uniquely in MGI or NovaSeq. (f) Percentage of reads with taxonomic and functional assignments per sample, presented as MGI to NovaSeq ratio. (g) A taxonomic overlap of matched samples, sorted by descending functional overlap in (c). (h) Mean Euclidean distance between functional profiles under simulated rarefaction, showing greater systematic distance between platforms than between technical replicates. (i) Recall comparison of the sequencing technologies, computed pairwise within matched samples using the union as a pseudo-reference.

The functional similarity (based on enzyme presence/absence) within matched pairs mirrored the taxonomic pattern, ranging from 80% to 97% (**Figure 5c**). Despite the high similarity, there were statistically significant differences in beta diversity between platforms (Euclidean distance PERMANOVA p<0.0001; Bray-Curtis distance PERMANOVA p<0.001). In pairs with lower functional overlap, MGI consistently yielded a higher number of detected enzymes per sample. Consequently, MGI systematically exhibited higher pathway recall compared to Illumina when assessed against a pseudo-reference (**Figure 5i**). A differential abundance analysis (MaAsLin2) revealed 241 enzymes significantly enriched in one platform (FDR-adjusted p<0.05). The bias showed a clear directional trend: 78% of the 100 most differentially abundant enzymes showed higher relative abundance in Illumina samples. The magnitude varied substantially. For example, trimethylamine-N-oxide reductase (X3.5.4.33) was significantly enriched in Illumina (coefficient=+0.38, q<0.001), while quinate/shikimate 5-dehydrogenase (X1.1.1.308) was severely underrepresented (coefficient=-1.22, q<0.001). Notably, these functional biases were most pronounced in central metabolic pathways, including carbohydrate metabolism, amino acid biosynthesis, and energy/cofactor metabolism.

To assess if the functional differences were caused by detection limits of rare functions, we examined the prevalence of the differentially abundant enzymes. We found the bias affected both ubiquitous and low-prevalence functions alike. For instance, many of the strongest discrepancies involved core functions present in of samples (e.g., X3.5.4.33 and X1.2.1.12), confirming the bias wasn’t due to sensitivity toward low-abundance features. Biases were also observed for sparser functions (e.g., serine endopeptidase, X3.4.21.19, present in 161 samples). This indicates a pervasive systematic shift affecting the entire functional landscape, extending beyond simple enzyme counts.

We hypothesized that sequence diversity underlies the observed functional discrepancies. To test this, a k-mer analysis was performed using the Jellyfish tool to evaluate sequence complexity. We found that MGI samples consistently contained 5–10 times more unique k-mers than their NovaSeq counterparts (**Figure 5d**). Despite this pronounced difference in k-mer diversity, the number of reads assigned taxonomic labels was similar across platforms. However, MGI samples yielded several-fold more functionally annotated reads (**Figure 5e**).This suggests that the increased sequence diversity in MGI data has limited impact on taxonomic profiling, but substantially improves functional annotation.

To investigate if functional differences were linked to taxonomic variation, we ordered sample pairs by decreasing taxonomic overlap and compared their functional similarity. No correlation was observed between the two (Spearman’s ρ = 0.005, p > 0.05, **Figures 5c** and **5g**), indicating that functional differences are not simply a consequence of taxonomic variation. Further analysis of genome coverage across a representative genomic fragment revealed platform-specific biases: MGI and Illumina datasets exhibited uneven and non-overlapping coverage patterns (**Supplementary Figure 5**). Specifically, regions highly covered by one platform were entirely missing in the other, which likely explains the functional discrepancies. While we observed no strong clustering by sequencing platform in a plot representing shared, MGI-unique, and Illumina-unique enzyme fractions, the values broadly overlapped, with MGI points consistently slightly higher (**Figure 5f**). This suggests the differences are not solely due to the sequencing technologies themselves, but are more likely linked to pre-sequencing protocols and uneven coverage.

To quantify differences between sequencing replicates and technologies, we simulated the effect of decreasing sequencing depth on the similarity of functional profiles. By iteratively subsampling reads and recalculating the Euclidean distance between profiles, we assessed the stability of intra- and inter-platform comparisons (**Figure 5h**). The results clearly demonstrate a systematic difference between the MGI and NovaSeq workflows, with the "MGI vs NovaSeq" comparison showing a consistently higher (approximately double) baseline distance than the technical replicate comparison ("MG1 vs MG2"). Importantly, this significant inter-platform distance persists even after rarefaction was applied to match the sequencing depth of the NovaSeq samples. This highlights that rarefaction alone is insufficient to fully mitigate systematic technological biases. Therefore, while technical replicates on the MGI platform are highly concordant, the choice of sample processing protocol introduces a more substantial source of variation.

## Discussion

Previous comparative analyses of MGI and Illumina sequencing technologies largely suggested their interchangeability for applications in metagenomics and clinical diagnostics (2),(14). Existing investigations, however, often suffer from limited sample sizes and fail to rigorously isolate the impact of the sequencing technology from other upstream methodological variables (e.g., library preparation or bioinformatics). Our study addresses these gaps by providing one of the first large-scale comparisons of MGI and Illumina NovaSeq sequencing technologies. We offer a comprehensive evaluation of MGI’s inherent robustness and consistency through extensive intra-platform comparisons and a direct, detailed assessment against the NovaSeq platform, aiming to clarify the reliability and comparability of these systems for advancing metagenomics.

Deeper analysis of sequencing quality revealed differences in GC content distribution and sequence duplication between the MGI and NovaSeq platforms, indicating potential sequencing biases. Published research has shown that GC content is a critical challenge, with workflows like MiSeq and NextSeq exhibiting major biases leading to falsely low coverage in GC-rich and especially GC-poor sequences (13). Despite these differences, our observed discrepancies, primarily within the 40-50% GC coverage range, were not systematic enough to influence major biological interpretations. We found strong agreement in taxonomic profiles, with over 94% overlap in species, genera, and families. Furthermore, key diversity metrics (Shannon entropy, UniFrac distances) and disease-microbiome associations did not differ significantly between the sequencing technologies. However, MGI samples tended to cluster more closely together than with their matched NovaSeq counterparts, suggesting that while intra-platform variation is minimal, inter-platform differences do exist.

Our observations revealed differences in the number and amount of enzymes detected in matched samples (the degree of functional similarity within some matched pairs was as low as 80% and differential enrichment analysis revealed 241 enzymes significantly more abundant in either platform). Overall, the functional overlap was considerably lower and did not align with the taxonomic deviations (Spearman correlation IZ > 0.05). This suggests that factors beyond mere taxonomic presence are influencing the functional profiles we observed. A recent study by the ABRF consortium sheds light on this issue, demonstrating how the initial amount of the starting material and subsequent PCR duplication rates can introduce significant biases (15). Their research, though focused on RNA-seq, found that low input material leads to a reduced number of detected genes and increased noise in expression counts. These conclusions are highly relevant to metagenomics, especially with the rise of higher multiplexing strategies.

Higher multiplexing, while cost-effective for large studies, reduces the input material per sample, often resulting in higher duplication rates and uneven coverage of the studied genomes. Taxonomic annotation tools are generally less affected by this uneven coverage because they use multiple genomic positions and rely on copy number for abundance determination. In contrast, functional profiling is much more sensitive; it depends on smaller genomic fragments and requires reasonably equal coverage across the genome, a condition difficult to maintain when input is limited and duplication is high. This supports our findings, where MGI samples, sequenced in smaller sets at higher depth, maintained higher and more consistent original coverage. Consistent with published work (16), which showed that low-prevalence functions are disproportionately lost at shallow sequencing depths, we found that although rarefaction was applied to normalize read depth, this correction was insufficient to eliminate disparity in functional detection.

## Conclusions

Our study confirms that MGI sequencing is a reliable and robust alternative to NovaSeq for taxonomic microbiome profiling, however, we observed differences at the functional level. Our findings underscore the critical need for methodological consistency when comparing results, particularly in studies utilizing a gene catalog.

For accurate and comparable outcomes, standardized sample preparation is a critical first step. Furthermore, experimental design should prioritize sequencing depth. We strongly recommend limiting the degree of sample multiplexing; this increases the amount of input material per sample in the prepared sequencing library, which is essential for minimizing coverage gaps and achieving a more uniform representation of the bacterial genomes. This comprehensive breadth of coverage is required for reliable functional annotations.

## Methods

### Cohort overview

The Estonian Microbiome Cohort (EstMB, n=2,509) was established in 2017 (12) as a part of the Estonian Biobank (EstBB), currently exceeding 202,282 genotyped adults across Estonia (17). The Estonian Microbiome Deep cohort (EstMB-deep) includes a subset of samples from the EstMB that have been deeply resequenced with MGISEQ-2000 platform (N = 1,729). Samples for EstMB-deep were randomly selected from EstMB.

### Microbiome sample collection and DNA extraction

Participants collected a fresh stool sample immediately after defecation using a sterile Pasteur pipette and placed it in a 15 mL conical polypropylene tube. They were instructed to collect the sample as close as possible to their study center visit. Samples were transported and stored at −80 °C until DNA extraction. The median time between sampling and freezing was 3 h 25 min (mean 4 h 34 min), and transport time was not significantly associated with alpha (Spearman correlation, p-value 0.949 for observed richness and 0.464 for Shannon index) or beta diversity p-value 0.061, R-squared 0.0005). Microbial DNA was extracted from 200 mg of stool using a QIAamp DNA Stool Mini Kit (Qiagen), following manufacturer’s instructions, after all samples were collected. DNA was quantified using a Qubit 2.0 Fluorometer with a dsDNA Assay Kit (Thermo Fisher Scientific).

### Metagenomic data processing

#### Illumina NovaSeq 6000 platform

The metagenomic paired-end sequencing was performed for whole EstMB cohort (N=2,509) by Novogene Bioinformatics Technology Co., Ltd. using the Illumina NovaSeq6000 platform, resulting in 4.62 ± 0.44 Gb of data per sample (insert size, 350 bp; read length, 2 x 250 bp). A NEBNext® Ultra™ DNA Library Prep Kit (NEB,USA) was used for generating sequencing libraries, according to the manufacturer’s recommendations. Sequencing reads’ quality control was performed using Fastqc (v 0.12.1) (18). Read trimming and adapter removal was performed using fastp (v0.23.2) (19) with default parameters before the rarefaction/subsampling step for both cohorts. The host reads were removed using Bowtie2 (v 0.6.5) (20) against the GRCH build 38, patch release (20),(21).

#### MGISEQ-2000 platform

Metagenomic paired-end sequencing was performed using DNBSEQI technology, resulting in 22.47 ± 4.2 Gb of data per sample (insert size, 350 bp; read length, 2 x 250 bp). Library preparation and circularization of equimolarly pooled libraries were done with MGIEasy FS DNA Library Prep Set (MGI Tech Co, Ltd, Shenzhen, China) according to the standard protocol. The sequencing was performed with MGISEQ-2000 High-throughput Sequencing Set (FCL PE150) according to the manufacturer’s instructions (MGI Tech, Shenzhen, China). Read quality control was performed using Fastqc (v 0.12.1) (18). Read trimming and adapter removal was performed using fastp (v0.23.2) (19) with default parameters before the rarefaction/subsampling step for both cohorts. The host were removed using Bowtie2 (v 0.6.5) (20) against the GRCH build 38, patch release (21).

### Bioinformatics and statistical analysis

#### Sample selection

For downstream analysis, we selected 1,729 samples sequenced with both platforms from the EstMB and EstMB-deep cohorts. To minimize bias, we first excluded sample pairs with fewer than 13 million or more than 20 million NovaSeq reads, as the number of NovaSeq reads was consistently lower than MGI (post QC). We then adjusted the number of MGI reads to match the NovaSeq read count within each sample pair using seqtk v1.3 (22). The final dataset consisted of 1,351 samples with approximately 20 million reads per sample. For the separate intra-sequencing comparison, we used the 53 samples sequenced twice with the MGI platform.

#### Read quality comparison

The FastQC files were aggregated and analysed using Python. The quality control was conducted to assess base sequence quality, adapter content, sequence length distribution, GC content, sequence duplication levels, and overrepresented sequences. Differences in GC content between sequencing platforms were evaluated using the Wilcoxon signed-rank test, and correlation was assessed using Pearson’s correlation coefficient. Samples exceeding sequence duplication thresholds - indicative of PCR amplification bias, low library complexity, or potential contamination - were flagged. Overrepresented sequences were identified based on FastQC’s criteria for sequence enrichment beyond expected distributions. For all downstream analyses, we included only sample pairs where both samples passed all quality criteria (N = 1,351).

#### Taxonomic profiling

Taxonomic profiling was done using Kraken2 v2.1.1 and Bracken v2.8 with a custom database (GTDB release 214) (23),(24). Kraken report files were processed using custom scripts (parse_kraken_NovaSeq.py, parse_kraken_mgi.py), retaining only taxa with relative abundance exceeding 0.01%. Relative abundance was defined based on the Kraken’s ’fraction_total_reads’ values and prevalence was calculated as the fraction of samples in which the species was detected.

## Statistical analysis

Differences in species counts between sample pairs were tested for normality using the Shapiro–Wilk test. As distributions deviated significantly from normality (p ≪ 0.05), comparisons between runs and platforms were performed using the Wilcoxon signed-rank test. Effect sizes were calculated using Cohen’s d.

## Further analyses

Alpha and beta diversity measures (UniFrac and procrustes analyses) were performed using QIIME 2 with default parameters (25). Functional predictions were made on a random subset of 700 samples with mi-faser with standard parameters (26). K-mers were calculated with jellyfish v.2.3.0 (27). Differential enrichment analysis was done with MaAsLin2 in its default mode (28). The association between centered log-ratio transformed species abundances and 33 common diseases (defined by ICD10 codes with cases in the EstMB cohort) was investigated using linear regression models. For each disease, the remaining samples served as controls. The models were adjusted for BMI, gender, and age, and the significance level was strictly controlled using a Bonferroni correction for the number of analysed species.

## Genome coverage

To evaluate sequencing coverage of key microbial species across samples, we selected four representative gut bacteria: *Akkermansia muciniphila*, *Bacteroides uniformis*, *Faecalibacterium prausnitzii*, and *Prevotella copri*. For each species, reference genomes were downloaded from the NCBI genome database (accessions: GCA_000020225.1, GCA_025147485.1, GCA_003324185.1, and GCA_025151535.1, respectively). Genome sequences in FASTA format were concatenated into a single reference file and indexed using Bowtie2 to create a searchable index (20).

To determine coverage, post-QC paired-end sequencing reads from each sample were individually aligned to the combined reference using Bowtie2. The alignment was converted to BAM format and sorted using SAMtools (29). Per-base genome coverage was then calculated with BEDTools’ *genomecov* function, using the reference genome file to define genome sizes (30). This process was repeated for each sample to quantify species-level coverage separately across all conditions.

Genomic coverage comparisons between the two sequencing methodologies were performed in Integrative Genomics Viewer (31). For illustrative purposes, a representative genomic fragment was randomly selected, clearly demonstrating the observed differences in coverage uniformity.

## Code availability

The source code for the analyses is available at GitHub: https://github.com/Chartiza/2024_Illumina_vs_BGI

## Acknowledgements

We express our gratitude to all individuals who made valuable contributions to the Estonian Microbiome cohort - Oliver Aasmets, Kertu Liis Krigul, Kreete Lüll and Steven Smith. We thank the Estonian Biobank research team: Andres Metspalu, Lili Milani, Tõnu Esko and Mait Metspalu for providing health data from Estonian Biobank participants. Data analysis was carried out in part in the High-Performance Computing Centre of University of Tartu, and we thank HPC Support Team of the Institute of Computer Science at the University of Tartu for delivering exceptional service and assistance in installing the necessary programs on the cluster. Special thanks to the Writing Retreat organized by the Institute of Genomics, University of Tartu for providing a conducive atmosphere for writing this paper. We also thank MMHP (Million Metagenomes of Human Project) for providing the sequencing for EstMB-deep cohort. Last but not least, we thank Łukasz Szydłowski for useful discussions during the early stages of this study and Rafał Mostowy for inviting Elin Org as a part of his EMBO Installation Grant, which established this project.

