## Supplementary Figures 1-9 for "A large-scale comparative metagenomic analysis of short-read sequencing platforms indicates high taxonomic concordance and functional analysis challenges"

Supplementary Information

### Kinga Zielińska, Kateryna Pantiukh, Paweł P Łabaj, Tomasz Kosciolek^#^, Elin Org^#^

Tomasz Kosciolek

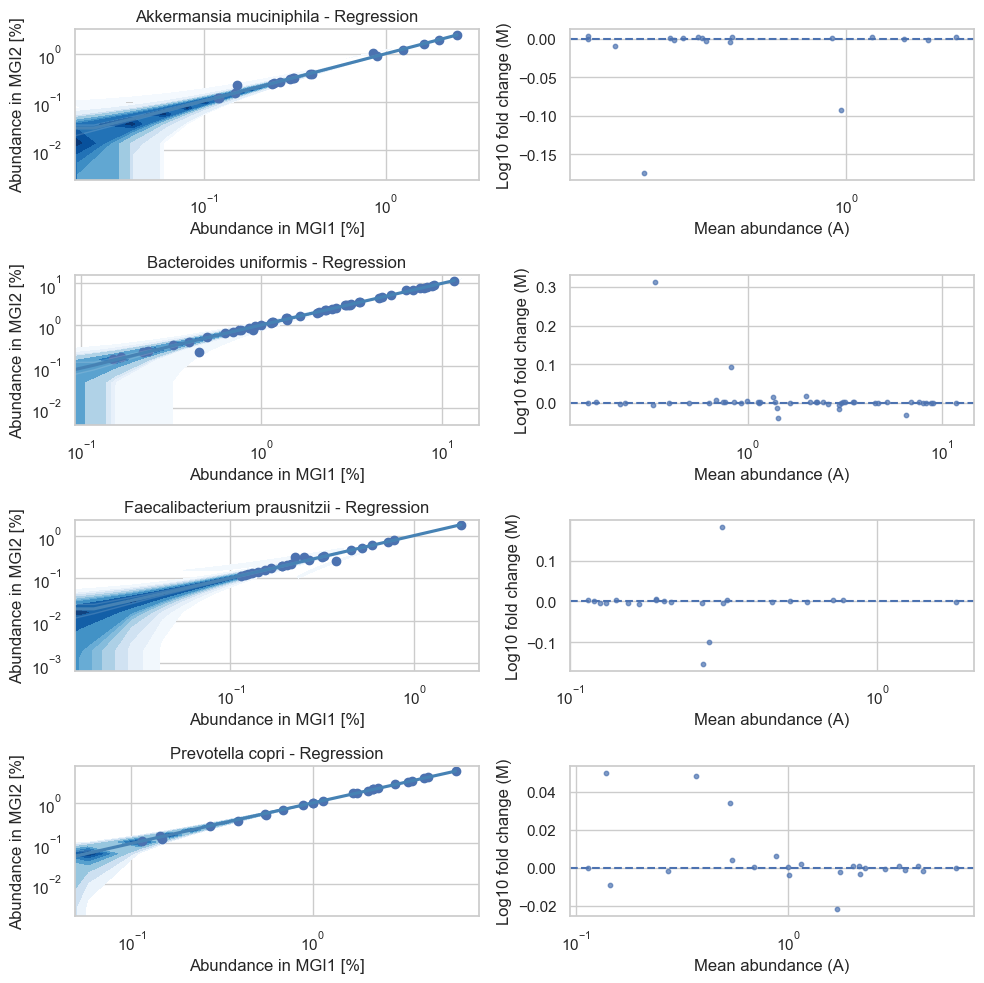

**Supplementary Figure 1.** Abundance correlations and residual plots for species of interest in the matching MGI-MGI pairs.

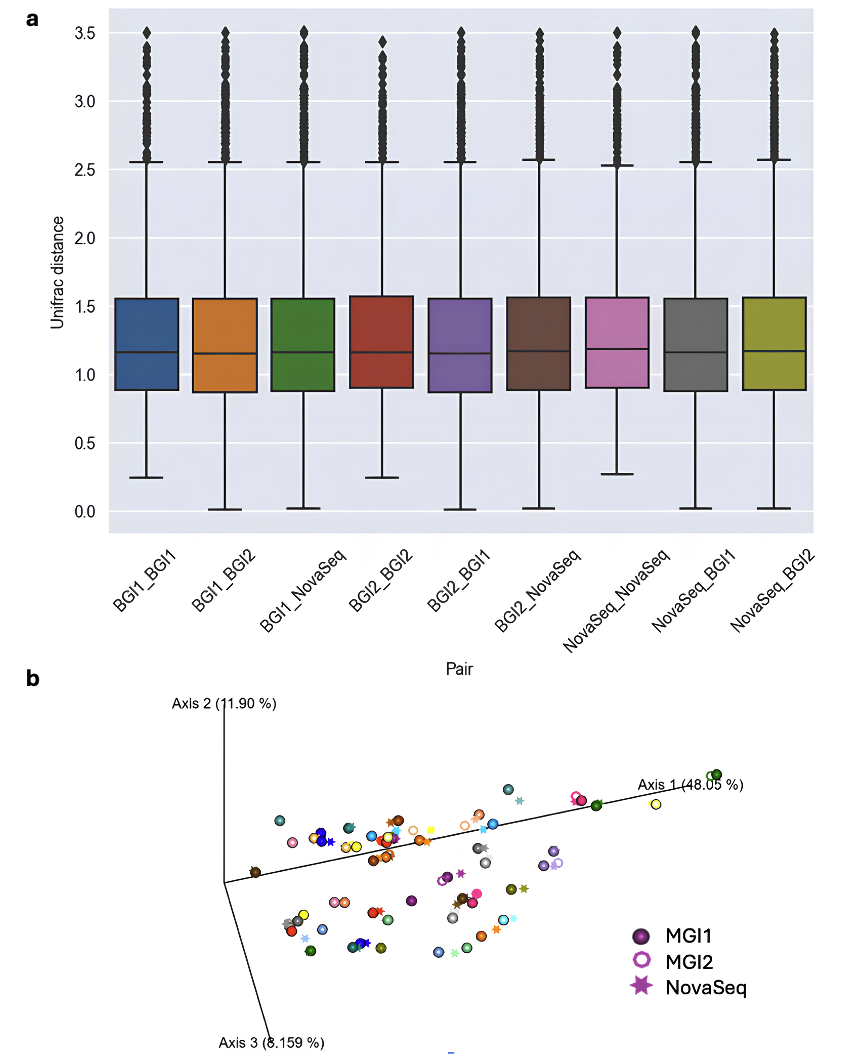

**Supplementary Figure 2.** Comparison of distances between MGI1, MGI2 and NovaSeq samples (N=53 samples). a) Shannon entropy; b) Beta diversity PCoA plot with Unifrac distance.

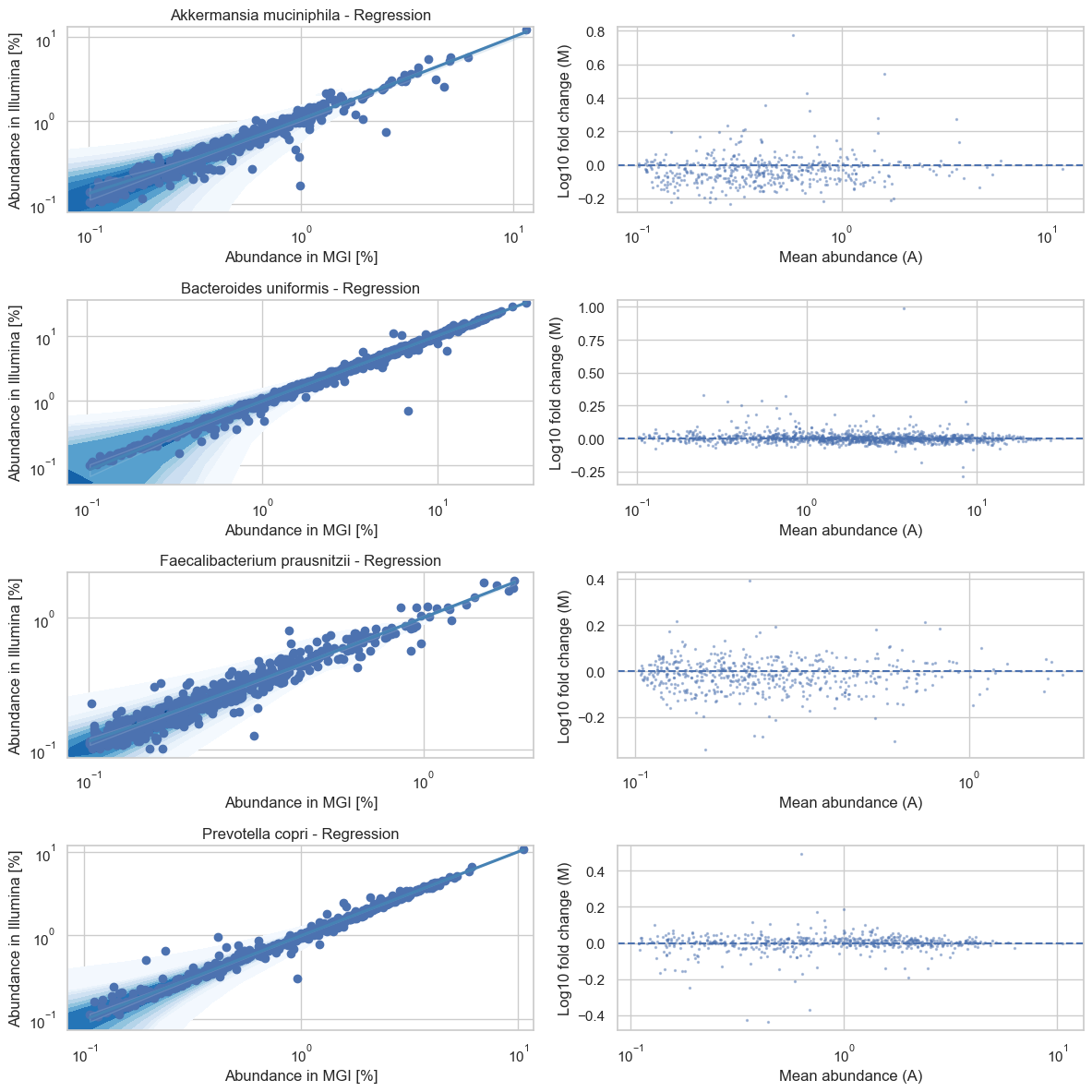

**Supplementary Figure 3**. Abundance correlations and residual plots for species of interest in the matching MGI technology and NovaSeq samples.

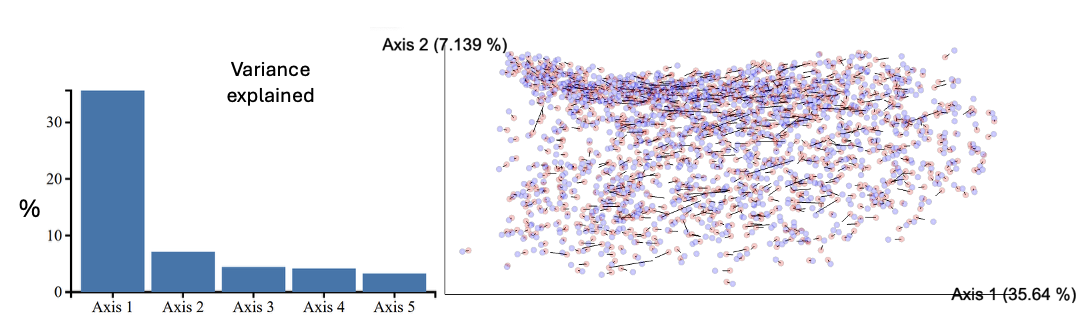

**Supplementary Figure 4**. Procrustes analysis with UniFrac distance: matched MGI technology and NovaSeq samples overlap well in the PCoA space.

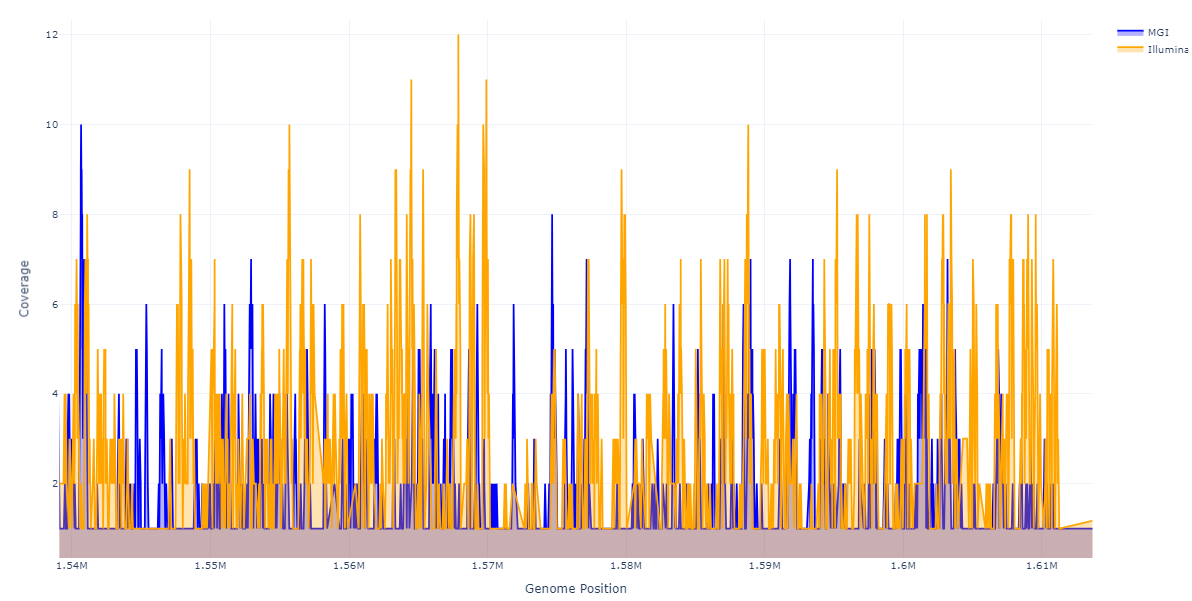

**Supplementary Figure 5.** MGI and NovaSeq coverage across an exemplary region of a bacterial genome considered in this study. A closer examination of the coverage revealed differing regions, explaining the observed functional discrepancies. The differences were not due to sequencing technologies, however, but were rather linked to pre-sequencing protocols and uneven coverage.

Supplementary results

#### Solving the mystery: matched MGI-Novaseq samples with a poor taxonomic overlap

While Figure 4d shows a very good overlap of species identified in matching MGI technology and Novaseq (Illumina) samples, the figure we obtained when we ran the comparison for the first time included a set of samples with a very poor overlap (Supplementary Figure 6a). The poor overlap was evident at different taxonomic levels: species, genus and family. Wondering whether the overlap could be a result of Kraken, a k-mer based taxonomy annotator used, we reran the annotation with reference-based mOTUs software. This was not the case, however, as we obtained a similar trend and a significantly different overlap between randomly selected well-overlapping and poorly-overlapping samples (p<0.001, Supplementary Figure 6b). The bimodal distribution was also not explained by any of the quality parameters, such as GC content, nor by the DNA concentration or date of sequencing. Furthermore, we confirmed that the poor overlap was not caused by random sequencing errors from the MGI technology. One sample, which was sequenced twice using MGI, showed a poor overlap with its Illumina counterpart but had a 92% overlap between the two MGI runs.

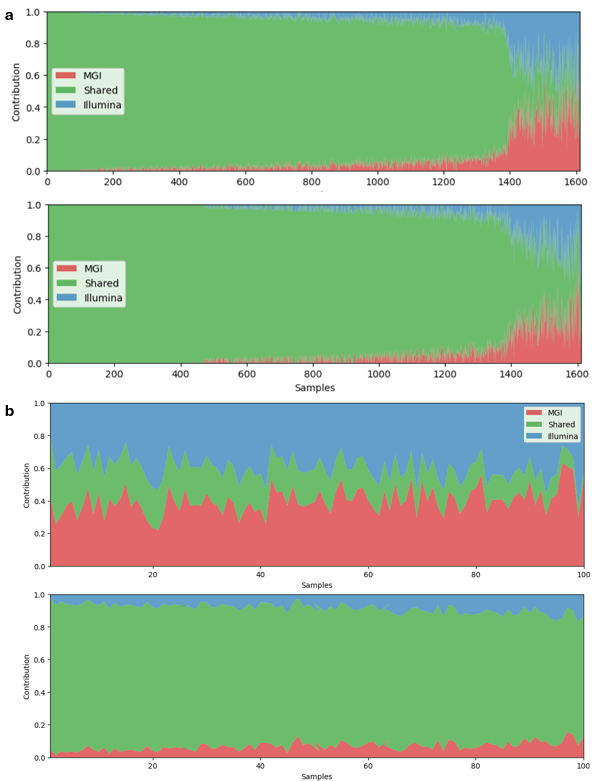

**Supplementary Figure 6**. (a) Overlap of genera (top) and families (bottom) in matched MGI technology and Illumina samples, where columns represent individual samples, produced by Kraken2+Bracken. Samples with a poor overlap (on the right side of the plots) appeared at different taxonomic levels. (b) Overlap of species in matched poorly-overlapping (top) and well-overlapping (bottom) MGI technology and Illumina samples, produced by mOTUs. The poorly-overlapping and well-overlapping subset was selected based on the samples in (a).

Our next step involved calculating beta diversity distances between the matched MGI and Illumina samples, and plotting them against the fractions of species shared by both (Supplementary Figure 7). We observed that pairs with a lower overlap were more distant from one another, regardless of the distance measure used.

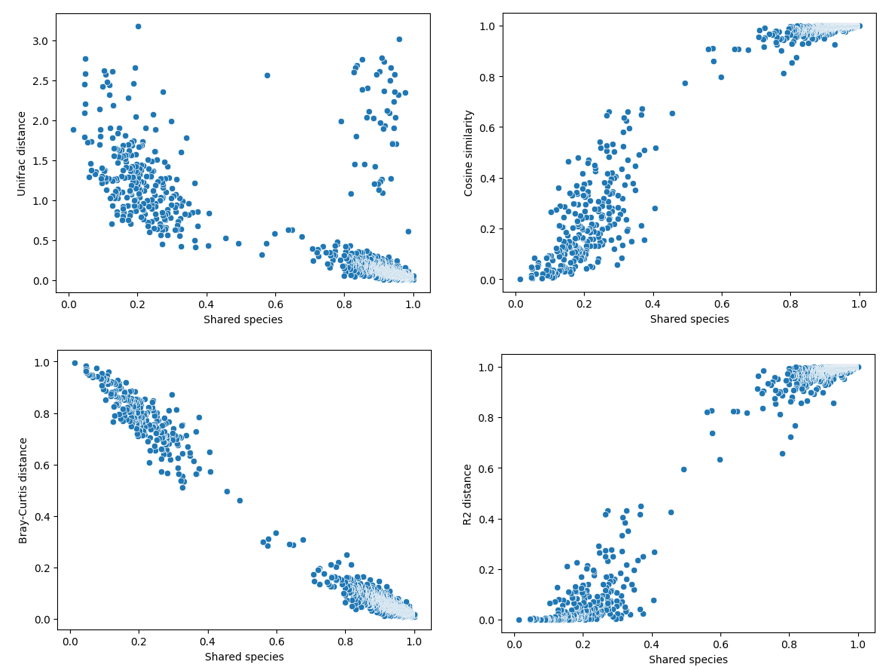

**Supplementary Figure 7.** Fractions of shared species versus distance measures for matched MGI technology-Illumina pairs.

Interestingly, we noticed that some Illumina samples would be substantially closer to a different MGI technology sample than to their original match. The newly located “match” would be the only sample very close to the original Illumina sample, while other samples would be located at a further, similar distance. This made us suspicious of potential mislabelling that could result in the poor overlap of the originally matched MGI technology-Illumina samples.

We made an attempt to “re-label” the poorly overlapping Illumina samples by matching them to MGI technology samples located at the closest distances, confirmed by multiple distance measures. The procedure allowed us to “re-match” 102 out of 261 poorly overlapping pairs. The resulting distances were of the expected range (Supplementary Figure 8a) and the taxonomic overlap reached the levels of the well-overlapping samples from Figure 4b (Supplementary Figure 8b). This confirmed our suspicions that the samples must have been mislabeled, which resulted in the poor overlap.

In our manuscript, only correctly matched samples were included. Any samples considered in this section, including samples which had been rematched, were not included in the main Illumina-MGI technology comparison.

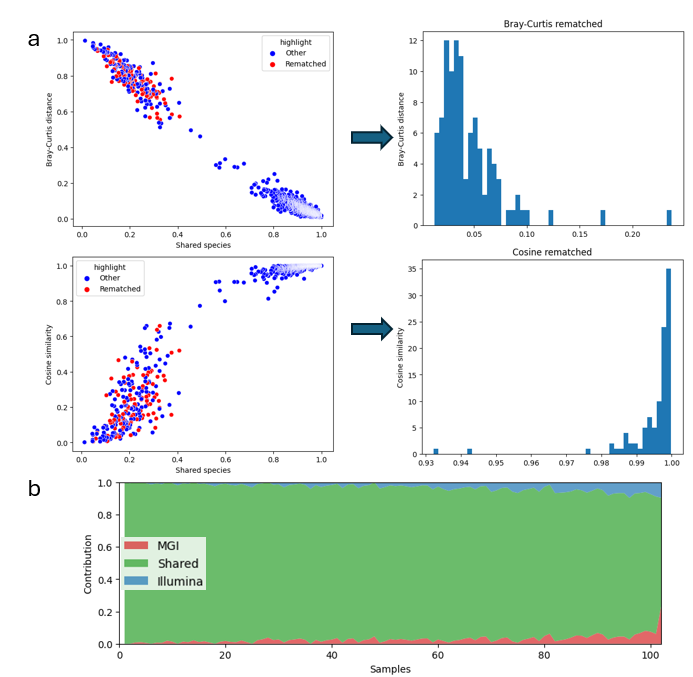

**Supplementary Figure 8.** (a) “Re-matching” of the poorly overlapping samples resulted in the new distances in the expected range. (b) The newly formed pairs displayed a good taxonomic overlap.

##### Cross-platform concordance in taxonomic profiles with Metaplan 4.1

In addition to Kraken2 + Bracken taxonomic profiling, we performed taxonomic comparisons for all sample pairs from the final dataset (N = 1,351) using the complementary marker-based method **MetaPhlAn v4.1**. Across both sequencing platforms, we identified a total of 2,466 species. Of these, 2.92% (72 species) were detected exclusively by one platform—53 species unique to MGI and 19 to NovaSeq (Supplementary Figure 9a). Most platform-specific taxa were rare, with a mean relative abundance of 3.55 × 10⁻⁶ %, markedly lower than that of shared species (4.16 × 10⁻² %) (Supplementary Figure 9b).

The majority of sample pairs exhibited high concordance, with over 90% of species shared between replicates (mean = 92.15 ± 3.83%) (Supplementary Figure 9c). Species prevalence and relative abundance were strongly correlated across sequencing runs, with R² values approaching 1.0 (Supplementary Figures 9d–9e). These values closely match those obtained using the k-mer–based Kraken2 + Bracken approach, indicating that the conclusions presented are robust and independent of the profiling method used.

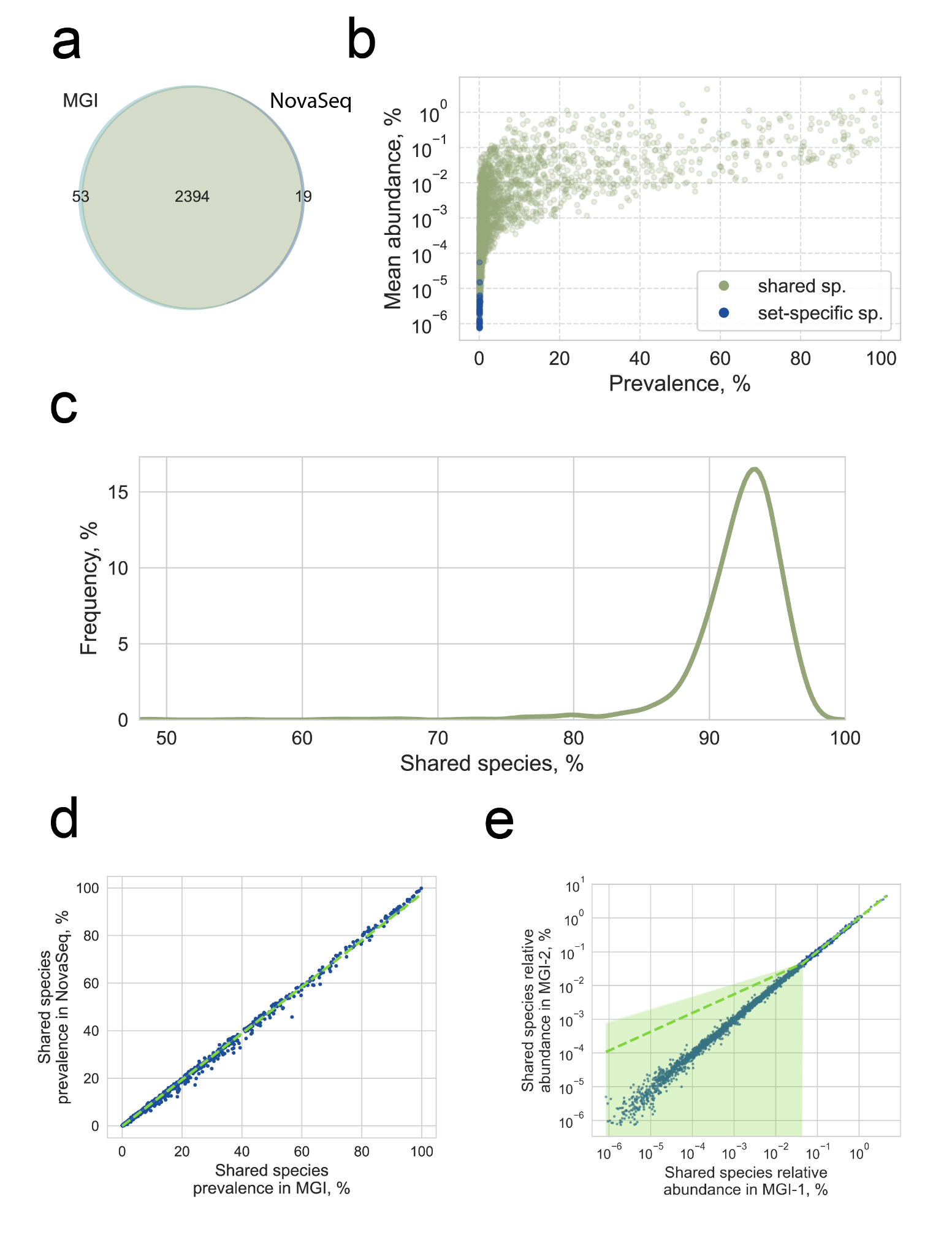

**Supplementary Figure 9.** Taxonomic composition comparison between sequencing platforms based on MetaPlan profiling (n=1,351) **a.** Proportion of species shared between platforms and those detected exclusively in MGI or NovaSeq. **b.** Prevalence and relative abundance of all detected species, with platform-specific species (found only in one platform) highlighted in blue. **c.** Distribution of the percentage of shared species within each sample pair. **d.** Species prevalence for taxa shared between platforms within sample pairs. **e.** Relative abundance of shared species within sample pairs.
